# Genomic Rearrangements Considered as Quantitative Traits

**DOI:** 10.1101/087387

**Authors:** Martha Imprialou, André Kahles, Joshua G. Steffen, Edward J. Osborne, Xiangchao Gan, Janne Lempe, Amarjit Bhomra, Eric Belfield, Anne Visscher, Robert Greenhalgh, Nicholas P Harberd, Richard Goram, Jotun Hein, Alexandre Robert-Seilaniantz, Jonathan Jones, Oliver Stegle, Paula Kover, Miltos Tsiantis, Magnus Nordborg, Gunnar Rätsch, Richard M. Clark, Richard Mott

**Affiliations:** Wellcome Trust Centre for Human Genetics, University of Oxford, OX3 7BN, UK; Memorial Sloan-Kettering Cancer Center, New York City, NY 10065, USA; Department of Biology, University of Utah, Salt Lake City, UT, 84112-0840, USA; Max Planck Institute for Plant Breeding Research, 50829 Köln, Germany; Department of Plant Sciences, University of Oxford, Oxford, OX1 3RB, UK; Department of Comparative Plant and Fungal Biology, Royal Botanic Gardens Kew, Ardingly, RH17 6TN, UK; John Innes Centre, Norwich, NR4 7UH, UK; Department of Statistics, University of Oxford, OX1 3TG, Oxford, UK; UMR INRA-Agrocampus Ouest-Université de Rennes 1, 35653 Le Rheu Cedex, France; The Sainsbury Laboratory, Norwich Research Park, Norwich NR4 7UH, UK.; European Bioinformatics Institute, Hinxton, Cambridge, CB10 1SD, UK; Dept of Biology and Biochemistry, University of Bath, Bath, BA2 7AY, UK; Gregor Mendel Institute of Molecular Plant Biology, Vienna, 1030, Austria; Center for Cell and Genome Science, University of Utah, Salt Lake City, UT, 84112-0840, USA; UCL Genetics Institute, University College London, WC1 6BT, UK

## Abstract

To understand the population genetics of structural variants (SVs), and their effects on phenotypes, we developed an approach to mapping SVs, particularly transpositions, segregating in a sequenced population, and which avoids calling SVs directly. The evidence for a potential SV at a locus is indicated by variation in the counts of short-reads that map anomalously to the locus. These SV traits are treated as quantitative traits and mapped genetically, analogously to a gene expression study. Association between an SV trait at one locus and genotypes at a distant locus indicate the origin and target of a transposition. Using ultra-low-coverage (0.3x) population sequence data from 488 recombinant inbred *Arabidopsis* genomes, we identified 6,502 segregating SVs. Remarkably, 25% of these were transpositions. Whilst many SVs cannot be delineated precisely, PCR validated 83% of 44 predicted transposition breakpoints. We show that specific SVs may be causative for quantitative trait loci for germination, fungal disease resistance and other phenotypes. Further we show that the phenotypic heritability attributable to sequence anomalies differs from, and in the case of time to germination and bolting, exceeds that due to standard genetic variation. Gene expression within SVs is also more likely to be silenced or dysregulated. This approach is generally applicable to large populations sequenced at low-coverage, and complements the prevalent strategy of SV discovery in fewer individuals sequenced at high coverage.

## Introduction

Whilst genome resequencing has become cheap and ubiquitous, and can readily determine variations such as SNPs and very small indels, the problem of identifying structural variants (SVs) and rearrangements remains a challenge, despite continual improvement in algorithms for calling SVs. The current gold standard for determining SVs between individuals is by *de-novo* assembly^1^. This requires very high-coverage paired-end sequence over a range of insert sizes, together with long-range information for scaffolding. Advances in long-read technologies^2,3^ are beginning to aid this process, but the relatively high cost and low throughput of this strategy limits its applicability to smaller numbers of genomes, and leaves open two important questions. First, whether an SV identified in an individual is unique, or is frequent enough to contribute appreciably to phenotypic heritability in a population. Second, whether an SV represents a local rearrangement, such as a deletion, inversion or tandem copy-number variant (CNV), or is long-range, such as a transposition^4,5^.

SVs are often revealed by the anomalous alignment of short-reads to the reference genome. Specific anomaly signatures characterize different types of SVs (Table 1). Thus, same-strand pairs indicate inversion, high read coverage duplications, abnormal insert sizes and unpaired reads are classified as indels.

**Table 1.** **(a)** MAGIC SV-QTLs classified by read pair anomaly type. SV-QTLs: total number of QTLs detected using each anomaly type (if the same QTL was detected by multiple anomalies then it is counted multiple times in this column), Unique: number of QTLs detected only by a single anomaly category, *cis*: number of *cis* SV-QTLs, *trans*: number of *trans* SV-QTLs. (**b)** MAGIC SV-QTLs classified by QTL type, after removing duplicates. SV-QTLs: number of structural variants of each type, *cis*: number of *cis* SV-QTLs (source and sink within 2Mb from each other), *trans*: number of *trans* SV-QTLS [Note that the total number of SV-QTLS is 10,275, of which 6,502 are distinct after removing overlapping events, and 5,899 unique to a single anomaly type.]

| trait type | SV-QTLs | Unique | *cis* | *trans* |
| --- | --- | --- | --- | --- |
| IP | 1997 | 833 | 1617 | 380 |
| ER | 184 | 165 | 112 | 72 |
| LIS | 2051 | 585 | 1677 | 374 |
| SS | 1950 | 1887 | 1358 | 592 |
| U | 2060 | 1998 | 1530 | 530 |
| U+LIS | 2033 | 431 | 1661 | 372 |
| **Total** | 10275 | 5899 | 7955 | 2320 |

**Table 1 (a)** MAGIC SV-QTLs classified by read pair anomaly type.
| SV type | SV-QTLs | *cis* | *trans* |
| --- | --- | --- | --- |
| duplication | 175 | 109 | 66 |
| indel | 3035 | 3035 | 0 |
| inversion | 1976 | 1373 | 603 |
| other | 1316 | 381 | 935 |
| **Total** | 6502 | 4898 | 1604 |

These signatures include excess read coverage (e.g., duplications, Copy Number Variants (CNV)), discordant distances between read pairs (e.g., indels) and inconsistent read orientation (e.g. inversions). These anomalies arise, often in combination, because the reads have been aligned to the wrong genome – the anomalies should disappear if instead the reads were aligned to the true genome. This idea is used by algorithms such as GATK^6^ and Platypus^7^ that identify small indels by local realignment, and in whole-genome reassembly by iterative realignment ^8^.

Many SV-calling algorithms utilize these read-anomaly signatures to identify SVs segregating in individuals sequenced at high coverage ^9–16^. These methods focus on short-range SVs because of the difficulties in distinguishing long-range rearrangements from read mapping errors. They are also designed to work best when calling SVs in individuals sequenced at intermediate to high coverage; for example, two of the most recent SV-callers, LUMPY^15^ and WHAM^16^ are most sensitive when sequence coverage is at least 10x. In other applications, e.g cancer resequencing, typical coverage is even higher, at 30x or above.

The problem of calling SVs from population sequence data presents additional challenges. Population studies are generally conducted for the purpose of genetic association, and consequently require large sample sizes. Population sequencing provides an alternative to genotyping by SNP arrays, simultaneously providing both haplotype reference panels for imputation^17^ and cohorts for disease mapping^18,19^. As the sample size increases, it becomes possible to reduce the coverage of each individual dramatically, yet still impute single nucleotide polymorphism (SNPs) accurately ^20^. Consequently one would want to be able to call SVs as well as SNPs and to test them for association. Although the information present in each sample is sparse, and therefore it would be difficult to call SVs (and SNPs) on an individual basis, by pooling information across samples it might be possible to determine common SVs analogously to the way SNPs are imputed.

A further challenge, which is not confined to low-coverage sequencing, is that presented by complex SVs. Unlike simple indels, inversions and transpositions, where a segment with well-defined breakpoints is affected, many SVs are composites of multiple events^21^, often driven by transposons and other repetitive mobile elements. Complex SVs resist simple classification, and it may be impossible to determine the precise sequence of mutations that occurred in the lineages separating the reference genome from that of the sequenced individual. Whilst current algorithms for calling SVs in simulated high-coverage human data can identify simple SVs with sensitivities of about 90% depending on the type of SV ^16^, they are less accurate when applied to real data, and their performance on complex SVs is unreported.

Nonetheless, even though it may be difficult to delineate complex SVs, there can still be strong evidence from read-mapping anomalies that an SV of some sort exists at a locus. If the intensity of its anomaly signature can be used as a proxy for the purpose of testing genetic association, then one need not delineate the SV precisely. It then follows that the genomewide information captured by these anomalies could be used to compute relationships between individuals based on their structural similarities alone, and hence to estimate the heritability attributable to this source of variation.

Here, we ask whether low-coverage population sequencing provides new ways for mapping SVs and estimating heritability, complementing the sequencing of fewer individuals at high coverage. As an illustration, we investigate the architecture and phenotypic impact of structural variation in *Arabidopsis thaliana*. Among natural accessions of Arabidopsis, structural variation is plentiful ^22^. The extent of rDNA repeats ^23^ and mobile transposable elements ^24^ vary between accessions, and variation in the overall amounts of both classes of repetitive sequence elements are complex traits, partially under genetic control. In this study we investigate all types of structural variation in Arabidopsis, including those not mediated by mobile elements. We show that long-range transpositions are common, and that structural variation has a significant impact on particular quantitative trait loci (QTLs) and on trait heritability, distinct from that explained by other types of sequence variation.

## Results

### Structural Variants as Quantitative Traits

We combined established ideas from signature-based SV identification with quantitative genetics to analyse structural variation in population sequence data. The following scenario motivates our reasoning: suppose an SV arose in a certain population ancestor, α, transposing a genomic segment *s* originating at a “source” locus *L* and targeting to a “sink” locus *M*. Source and sink can be coincident or unlinked, but for the moment, suppose they are unlinked. If the event is transposon-mediated, then the segment *s* is duplicated to *s’* at *M*, and possibly altered, leaving the original *s* at *L*. Once random chromosomal assortment and recombination has occurred, in the present-day population there will be a mix of individuals carrying the segment at neither, one or both loci.

In the descendent population, one individual is sequenced and becomes the reference genome. Depending on the choice of reference individual and the mechanism of transposition, the reference might carry zero or one copies of *s* at the source and of *s’* at the sink.

Assume the reference has one copy of *s* and zero copies of *s’*. A population sample will contain individuals with mix of all possible configurations at source and sink. But only individuals that inherited the haplotype descended from α at the sink carry the transposed segment, regardless of their haplotype at the source. The individuals are sequenced with short-reads, and the reads are mapped to the reference genome. Individuals carrying the transposition *s’* at the sink will have reads spanning the breakpoint that split between source and sink. Hence read mapping anomalies apparently originating at the source will be enriched in those individuals carrying the sink haplotype α: genotypes that tag α at the sink will be associated with anomalies at the source.

If on the other hand the reference contains both *s* at the source and *s’* at the sink then those individuals that did not inherit the haplotype α at the sink will appear to carry a deletion there. Reads with anomalously large insert sizes will map to the sink and will be associated with genotypes tagging the haplotype α at the sink – the generative role played by the source will be invisible.

Similarly, by considering situations where the source and sink are coincident – for example tandem duplications – in a population we would expect to encounter a mix of short-range *cis* and long-range *trans* associations between different classes of read-mapping anomalies and genotypes, depending on the diverse histories of each structural variants.

To apply these ideas in practice, we count the numbers of anomalous reads mapping to each source *L* in a population sample, treat it as a quantitative trait, and proceed to identify genetic loci containing variation that correlate with variation in the trait. This procedure defines a SV quantitative trait locus (SV-QTL) linking *T_Li_*, the number of anomalous reads mapping to locus *L* in individual *i* and the haplotype *H_Mi_* at sink locus *M* in individual *i* (Figure 1, **Methods**). *cis* SV-QTLs where the source and sink overlap indicate local structural variants such as CNVs, deletions and inversions; *trans* SV-QTLs indicate transpositions (insertional translocations) or larger scale rearrangements. In this way we can determine whether an SV is in *trans*, its originating haplotype, which individuals now carry it (**Figure S1**), and its frequency (**Figure S2**).

**Figure 1.**
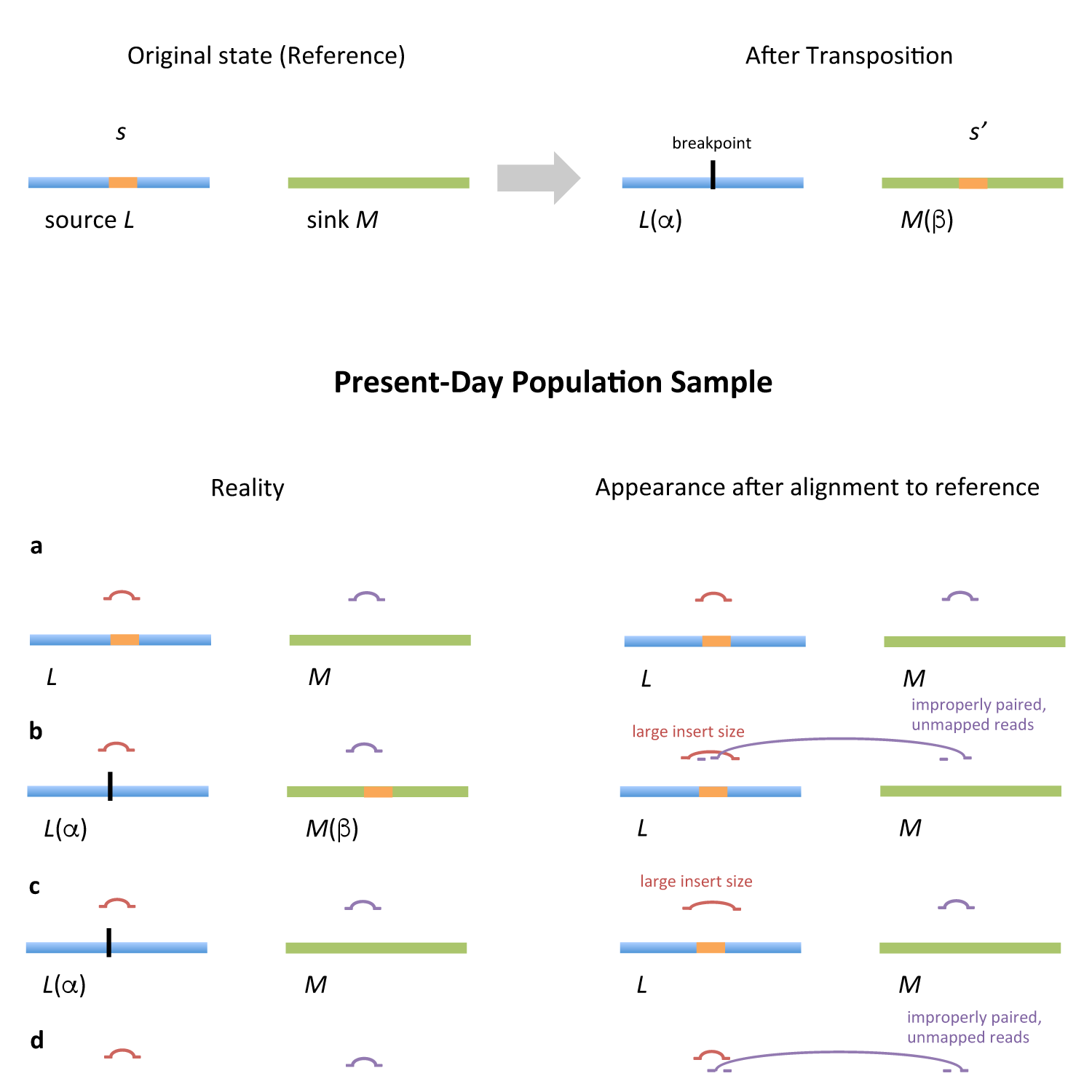
Effects of a transposition on short-read mapping. Chromosomes are horizontal bars and read pairs are pairs of horizontal lines linked by curves. Upper shows a population ancestor corresponding to the reference genome (left) undergoing a transposition (right), in which a segment *s* at source locus *L* with haplotype context *α* is copied to *s*’ at recipient sink locus *M* with haplotype context *β*. Lower shows all four possible combinations (a-d) of source *L* and sink *M* haplotype in descendants. On left are shown the alignment of reads to the true haplotypes, where there are no read-mapping anomalies. On right are shown the various read-mapping anomalies that arise, depending on the true haplotype backgrounds at source and sink, upon alignment to the reference genome.

We interpret the matrix of SV-traits across all loci as a Euclidean representation of haplotype space, in the sense that, if two individuals are genetically similar then their SV-trait vectors should be close together. Consequently we define a genome-wide similarity between individuals based on the similarity of their anomalies, as a weighted average of their locus-specific similarities. Taken across all individuals, these generate a structural variation similarity matrix, analogous to a SNP-based genetic relationship matrix. This matrix was used to estimate the heritability of a phenotype with respect to structural variation, and compared to the heritability associated with SNP variation.

### Structural Variation in Arabidopsis

We used our strategy to map *cis* and *trans* SVs in the 120Mb genome of the plant *Arabidopsis thaliana.* We sequenced 488 of the *Arabidopsis* Multiparent Advanced Generation Inter-Cross (MAGIC) recombinant inbred lines^25^ at ~0.3x coverage using 51bp paired-end Illumina reads. The MAGIC lines descend from 19 ancestral founder accessions that have been sequenced at high coverage^8^ (**Table S1**) such that each ~120Mb genome is a mosaic of the 19 founder haplotypes. Consequently we expect most SVs segregating in MAGIC to also segregate in the founders, thereby providing a means to verify any SVs we detect. The choice of MAGIC lines rather than natural accessions means that the confounding effects of population structure and of selection are largely absent from the population. Very rare alleles with frequency below 1/19=4.5% are uncommon, increasing the power to detect QTLs. However, MAGIC QTL mapping resolution is also poorer, at ~200kb, compared to ~10kb in natural accessions.

We mapped the reads to the TAIR10 reference using Stampy ^26,27^ and inferred the mosaic of each line using a hidden Markov model (HMM) implemented in the software ‘reconstruction’ available on request from the authors. The algorithm uses as input SNP calls for each MAGIC genome, and the set of of 1.2M biallelic variants in the 19 founders (excluding loci tagged as within transposons, and those sites called as heterozygous or multi-allelic in the founders)^17^, and finds the most likely sequence of haplotype assignments for each chromosome. Because the lines were called at low coverage, most SNP sites were not covered by reads in an given; consequently we called on average 301k SNPs per line (using GATK^6^) (ie a randomly sampled of ~25% of the 1.2M sites). However, this amount of data is sufficient for the HMM to determine the founder mosaic accurately; we estimated by simulation that the algorithm can delineate the mosaic breakpoints (which correspond to recombination events) to within ~2kb (data not shown).

Using this procedure, we reconstructed each MAGIC genome into ~34 haplotype blocks on average with mean size 3.48Mb, representing contributions from about 11 founder haplotypes (**Table S2**), and imputed the full variant catalogue into each lines. Comparison of imputed SNPs with 782 GoldenGate SNP genotypes measured in 370 of the MAGIC lines ^25^ showed 98% concordance.

To map SVs, we divided the reference genome into 11,915 abutting source loci, each 10kb wide, and computed six measures of anomalous read mapping in each locus (6*11,915 = 71,490 SV trait vectors) (**Methods**, Table 1a, **Table S3**). Four of these measures address different types of anomalous read mapping that provide evidence of specific anomalies, namely high read coverage for duplications, strandedness of reads for inversions, anomalously large insert size for translocations and unpaired reads for deletions. The remaining two measures are linear combinations of other measures that could co-exist.

Genetic association between each of the SV-trait vectors and the local haplotype space was determined using a one-way ANOVA. We chose to determine association at the level of haplotypes rather than SNPs for two reasons. First, the founder haplotype space in the MAGIC lines is well-defined, and measuring association with haplotypes can capture relationships invisible at the level of SNPs. Second, the set of haplotype tests - defined by the union of all the breakpoints, comprising 16,700 haplotype blocks, such that the ancestral haplotype of all lines is unchanged within each block – means about 75 times fewer tests are performed, thereby speeding up the procedure (**Methods**). To determine genome-wide significance thresholds for SV-QTLs we performed 100 phenotype permutations for each trait and then fitted extreme value distributions (evd) to the genome-wide maxima of the permutations (**Methods**). We merged together probable duplicate overlapping SV-QTLs identified by multiple anomaly types.

After removing duplicates we identified 6,502 SV-QTLs at 1% study-wide false discovery rate (evd P<0.001) (**Table S3**). Of these, 1,604 (25%) were *trans*, defined as mapping over 2Mb from the source. Overall, 4,073/11,915 (34.2%) source loci harboured structural variants. Whilst we have greater power to detect larger SVs, 2,379 overlapped annotated indels shorter than 2kb^8^.

The likelihood that a structural trait vector has an SV-QTL increases with its variance (Figure 2). SV-QTLs are enriched around centromeres, as expected. Away from the centromeres, Figure 2 also shows that bins with variable SV traits are isolated, rather than in clusters. Figure 3a shows the genome-wide distribution of SVs segregating in one MAGIC founder, Ler-0. Figure 3 and **Table S3** show trans SV-QTLs link all five chromosomes.

**Figure 2.**
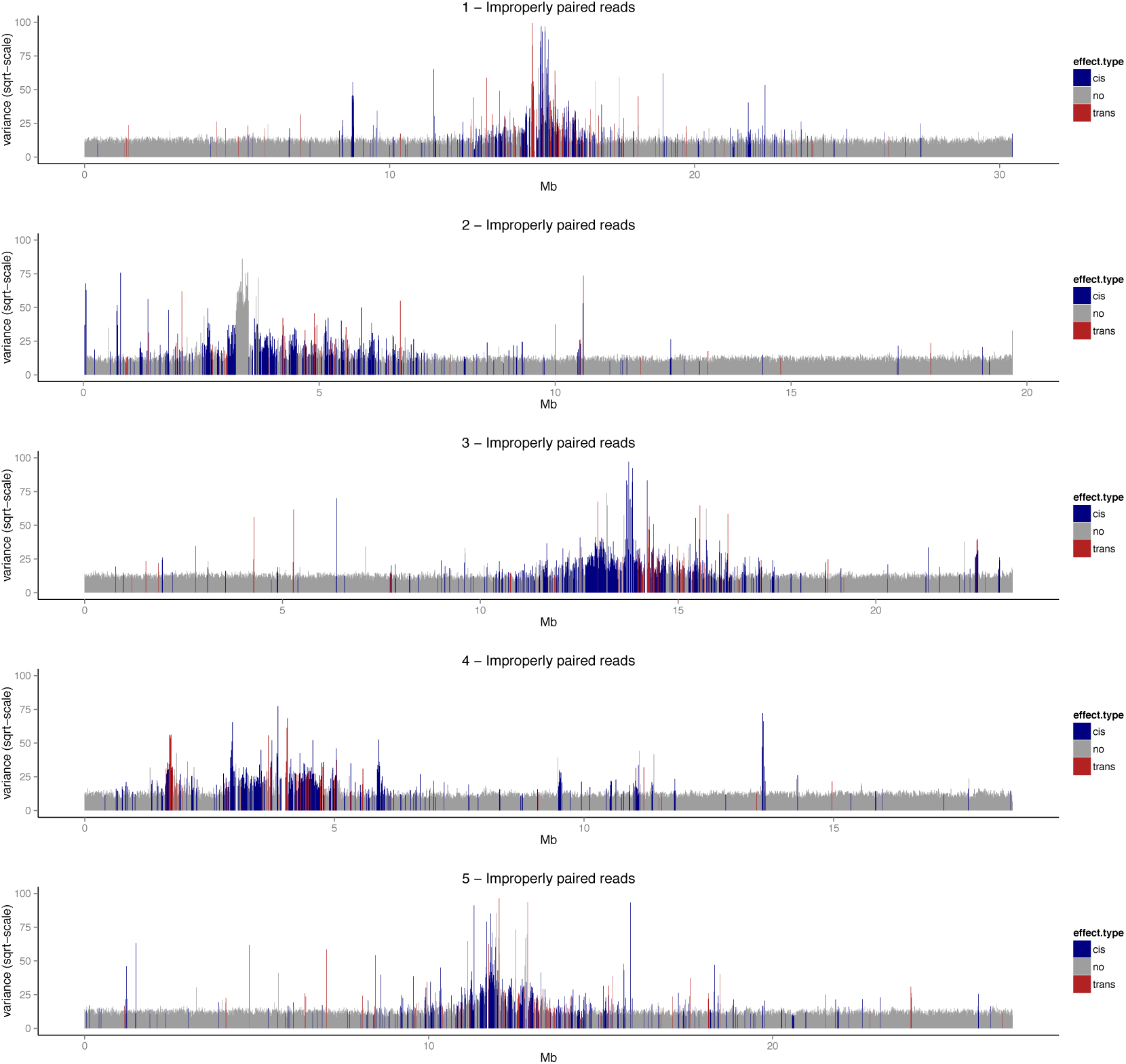
Genome-wide distribution of the variance for the trait “improperly-paired reads” (the number of reads mapping to a locus with mapping anomalies), computed in 10-kb windows. The x-axis shows genomic position and the y-axis the variance of each trait vector scaled by its mean. Each vertical line corresponds to a window. Those with SV-QTLs are blue (*cis*) and red (*trans*). Centromeres are marked by pink bars.

**Figure 3.**
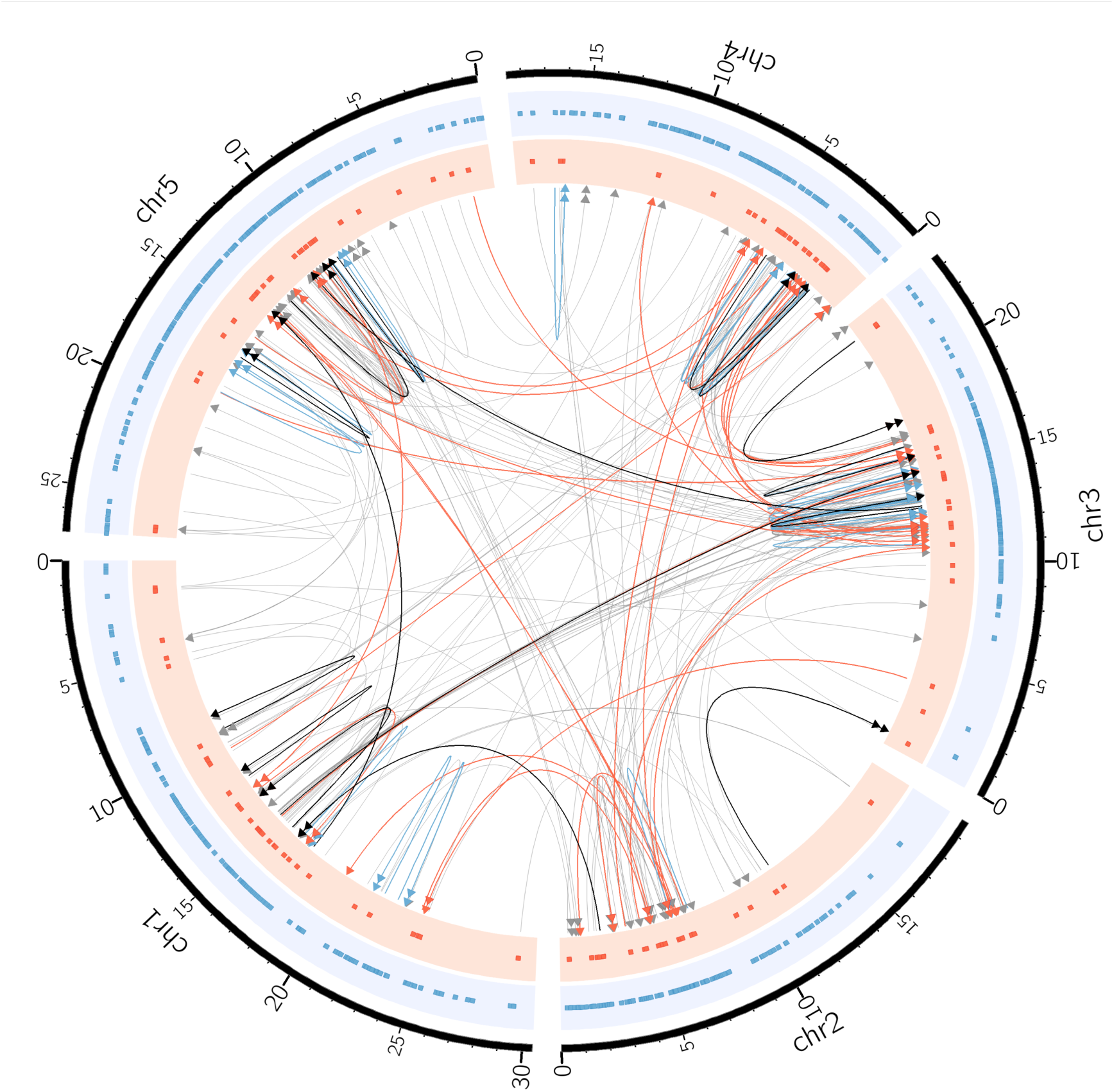
Structural variants segregating in the accession Ler-0. The grey directed lines show SV-QTLs with the arrows pointing towards the sink locus. Red and blue links indicate 37 *trans* and 30 *cis* SV-QTLs confirmed by *de novo* contigs. The black links show 16 SVs confirmed by PCR (7 *cis*, 9 *trans*). Double arrows in links indicate inversions. The dots in the red and blue tracks mark the sources (*trans* and *cis*, respectively) of all SVs associated with the Ler-0 haplotype.

In 319 SVs we were able to pinpoint both breakpoints, using contigs from *denovo* assemblies of the 19 founder genomes^8^ (see validation section below). Mean SV size was 53kb in these SVs, and the largest was 189kb. Thus the many of the SVs we discovered are too large to be due to insertions of small transposable elements. This probably reflects our lack of power to detect very small events, but also emphasizes that not all SVs are driven by mobile elements.

#### Validation

Genome-wide confirmation of SVs using short-read sequence is challenging because SV breakpoints often associate with transposons and repeats that hinder read-mapping and reassembly. However, among our SV-QTLs are several known rearrangements. These include trans SV-QTLs linking a cluster of rDNA repeats at ~14.2Mb on chromosome three to clusters at the ends of chromosome two. Polymorphisms in these clusters are implicated in massive genome size variations among *Arabidopsis* accessions^28^. We also identified the known knob inversion on chromosome 4 as reciprocal transpositions linking 1.61Mb and 2.65Mb^29^, and a 93kb inverted transposition identified previously in a cross between Ler-0 and Col-0^30^, and found it was present in 12 MAGIC founders.

To validate further SVs we compared our SV calls for the founder accession Ler-0 against two Ler-0 contigs (chr3:16.65-17.02Mb, chr5:25.06-25.23Mb) that were independently re-sequenced and manually reassembled^31^, thereby constituting a gold standard for comparison. The chromosome 3 contig (Figure 4) is enriched in SVs (83 indels, 31 larger than 100bp), consistent with our analysis: 42 SV-QTL sources (36 *cis* and 6 *trans*) are in this region and 4 *trans* SV-QTLs map into it. As would be expected, the sources of these SV-QTLs are within gaps in the contig. Furthermore, alignment revealed two long-range SVs within the contig (a transposition and a duplication which align to chromosomes four and two, respectively), which coincide with the source and sink of two *trans* SV-QTLs mapped within the contig. Similarly, in the chromosome 5 contig, 6 *cis* SV-QTLs correspond to deletions (**Figure S3)**.

**Figure 4.**
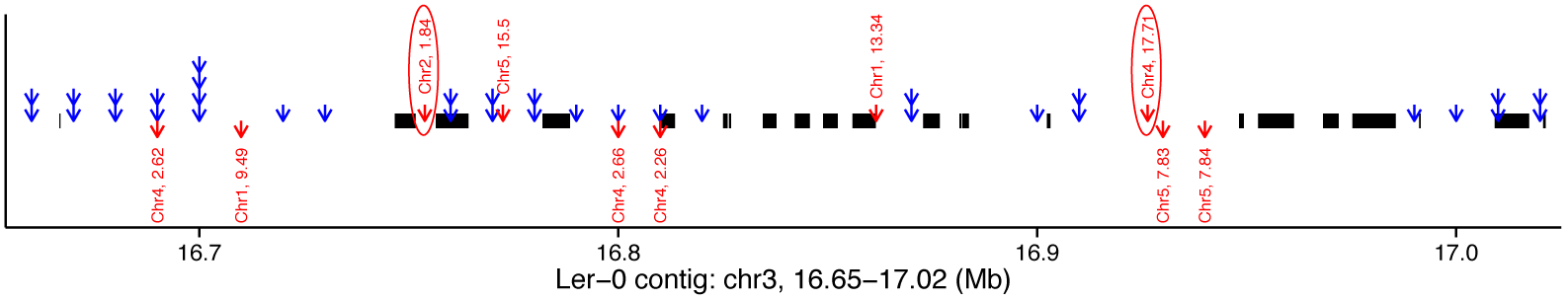
Alignment of a manually assembled contig from Ler-0, chr3:16.65-17.02Mb to the reference annotated with SV-QTLs. Thick black lines show alignments to reference genome. Blue arrows show the sources of *cis* SV-QTLs; stacked arrows mean multiple read anomaly traits had SV-QTLs. Red arrows display *trans* QTLs with arrows starting from the source and pointing towards the sink. Gaps in the contig alignment indicate loci where Ler-0 did not align to the reference, with the exception of two transposed segments that mapped to chromosomes 2 and 4 at positions concordant with the sources of two *trans* SV-QTLs (circled).

We also used an independent *de-novo* assembly of Ler-0 built from long PAC-BIO reads, Genbank accession GCA_000835945.1^32^ to validate our trans SV predictions. This assembly was constructed algorithmically without manual revisions, and so is not guaranteed to be correct. Further, the Ler-0 individual sequenced in the PAC-BIO assembly was different from the individual that founded the MAGIC population and therefore might carry private structural variations. Nonetheless, we expect it to be more accurate and contiguous than a Ler-0 assembly built from short Illumina reads alone. We took those 3080 Illumina paired-end reads for Ler-0 from ^8^ that carried large insert size mapping anomalies when mapped to TAIR10 and that mapped to the sources of our predicted Ler-0 *trans* SV-QTLS, and then mapped them to the PAC-BIO assembly using bwa^26^. These Illumina reads are from an individual grown from the same batch of seeds used to found the MAGIC population in ~2007, and should therefore share the same structural variants. Read anomalies that gave rise to correct SV predictions should map contiguously to the PAC-BIO assembly, under the assumption that the latter assembly is a more accurate representation of the Ler-0 genome. We found 2460 (80%) of these formerly split Illumina read pairs now mapped contiguously, defined as both members of a read-pair mapping to the PAC-BIO assembly with an insert size below 600 bp.

With the exception of these manually assembled Ler-0 contigs and the provisional Ler-0 PAC-BIO assembly, the MAGIC founders are not contiguously reassembled into a genome-wide gold standard reference panel. Nevertheless, they provide some information to test our SV predictions. To do this, at each SV-QTL we predicted which founder haplotypes carried SVs at the origination of the population. Using the low coverage data for the 488 MAGIC lines, at each SV-QTL we predicted which group of founders carried the SV allele *vs* the reference allele based on correspondence between their SV-trait value and predicted founder allele, using the fact that SV haplotypes will have elevated anomalous reads at the source. We were able to do this confidently at 2,391 SVs where the founders divided into two groups, the remainder having complex multi-allelic SV predictions (**Methods**). We then examined the independently-collected high-coverage reads in each the 19 MAGIC founders ^8^ for read-mapping signatures that supported the predicted grouping of founders at each SV. We counted the read pairs linking source and sink at each of the 2,391 SVs in the 19 high coverage founders. At 1,585/2,391 (66.3%, FDR 7.5%) SVs we observed significant differences in anomalies between the predicted groupings of founders (**Figure S4**, which also shows that the majority of SVs were mapped within 50kb). In the founders, the mean SV allele frequency was 6/19=31%. Only 387 (12%) were private to a single founder (**Figure S2**), in contrast to the fraction of SNPs (45%) that are private to a single founder^8^.

A related analysis using low-coverage reads from the 488 MAGIC genomes, but independent of founder predictions, (**Methods**) supported 1228/2391 (51.3%, most also supported by the founder genomes) and 1631/4111 (39.7%) of those remaining SVs without founder predictions. In total 2,965/6,502 (45.6%) SVs were supported by either method.

#### Breakpoint Prediction and Confirmation

In order to estimate SV sizes and identify SV breakpoints that could be tested experimentally by PCR, we next assembled the high-coverage sequence data for the MAGIC founders into *de-novo* contigs. No scaffolding was attempted in order to produce conservative high-quality short contigs, each up to a few kilobases long. We aligned these contigs to the reference to find alignments split between sources and sinks. We mapped breakpoints for 420 SV-QTLs (**Methods, Table S4**): in 319 SV-QTLs both breakpoints were identified. We designed PCR primers around 77 breakpoints from 45 SVs (both breakpoints in 7 SVs, and one in each of the remaining 38). We validated 37 SVs (83%), comprising 61 (79%) breakpoints, in 14 *cis* (6 inversions, 7 transpositions, 1 indel) and 23 *trans* (23 transpositions, 13 with inversions) SV-QTLs (**Table S5**).

Consistent with our difficulties in predicting biallelic founder alleles, in 11 SVs the breakpoints were polymorphic among the founders carrying the SV, and in 5 transpositions the orientation of the SV differed between founders. These results emphasise the difficulties frequently reported when delineating SVs.

#### Effects of SVs on phenotypic QTLs and gene expression

We next investigated associations between SVs and 9 physiological phenotypes, either previously published ^25,33^ or new to this study (**Table S6**). We found 16 distinct SV-QTLs (8 in *trans*, **Table S7**) that overlap physiological QTLs. In some cases, regressing the SV-trait from the physiological trait ablated the physiological QTL, consistent with, albeit not proof, that the SV is causal. This is illustrated by a QTL for germination time^25^ on chromosome 3, which is ablated by a *cis* SV-QTL for unpaired reads at around 15,936,650-15,951,640bp (Figure 5a,b). Our analysis predicted that 7 founders would carry a deletion at this locus, which was confirmed by the independent founder sequences (Figure 5c), revealing a 15kb deletion of three genes, *AT3G44240* (Polynucleotidyl transferase, ribonuclease H-like superfamily protein), *AT3G44245* (pseudogene of cytochrome P450, family 71, subfamily B, polypeptide 21), and *CYP71B38* (*AT3G44250*, cytochrome P450, family 71, subfamily B, polypeptide 38). Other SVs segregate nearby, but with allelic patterns inconsistent with the trait and therefore unlikely to be causal. It is therefore probable that the causal variant(s) lies within the deleted region. The three genes are not known to affect germination, although a mutant of another Polynucleotidyl transferase, *AHG2* (*AT1G55870*) does ^34^.

**Figure 5.**
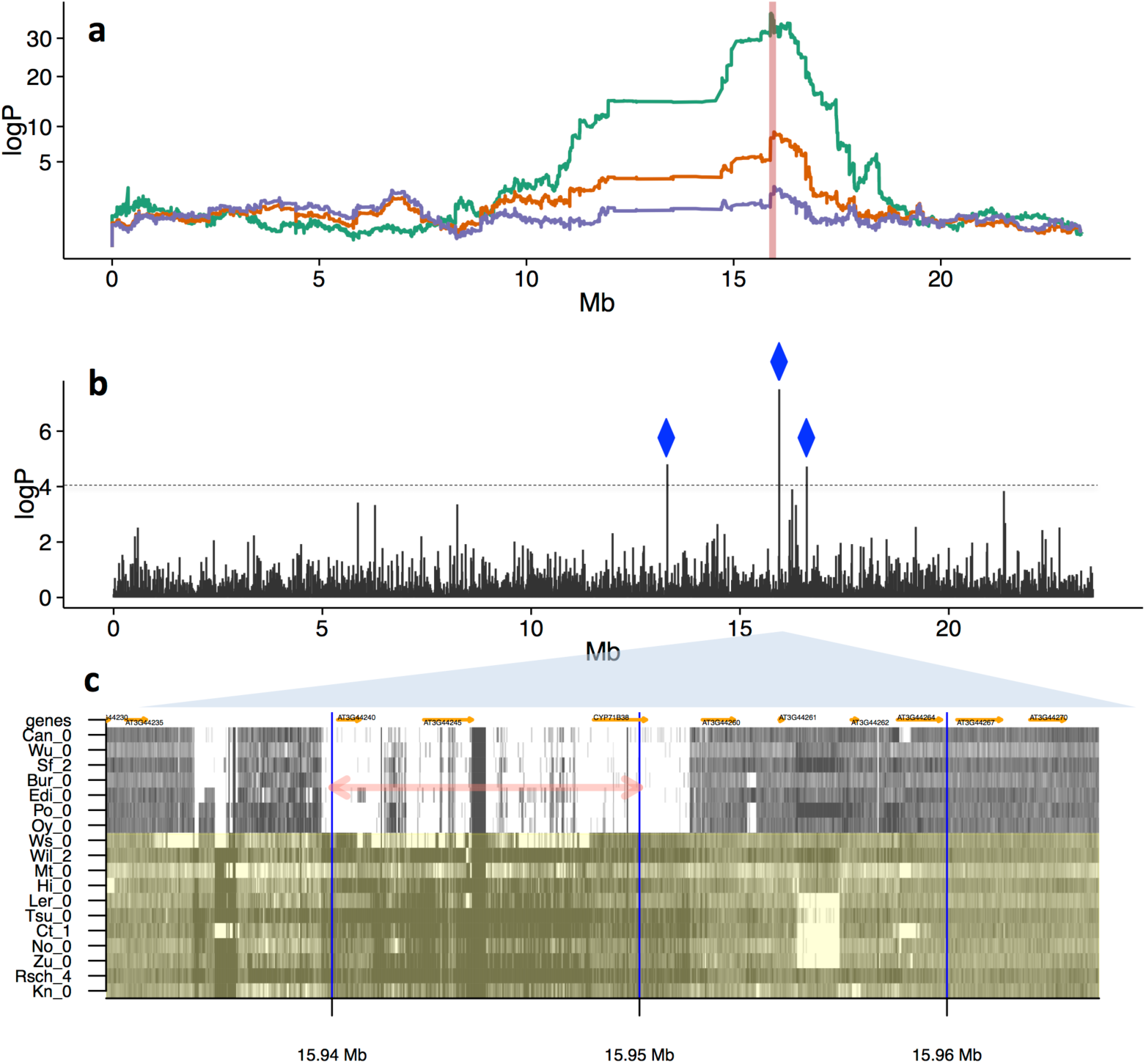
Association of haplotypes and SVs **(a)** Genome scans over chromosome 3 (x-axis: genomic position, y-axis: logP of association). Orange: association of local haplotype with germination time (days), peaking at 15.93Mb. Green: association of local haplotype with the SV trait unpaired reads at the source locus 15.94-15.95Mb (indicated by the vertical red line), explaining 8.13% of the variance in germination time, with an SV-QTL mapped at the same position as the germination QTL. Purple: residuals of germination time after regressing out the SV trait, ablating the QTL. **(b)** Chromosome-wide Pearson correlations between germination time and the numbers of unpaired reads measured at each 10kb source locus (x-axis: genomic position, y-axis: -log_10_ P-value of test that the correlation is zero). Three source loci correlate strongly with germination (logP>4), all with *cis* SV-QTLs (blue diamonds). **(c)** Structural variation in the MAGIC founders. Shown is the read coverage in 18 accessions (labelled on y-axis), over ~30kb surrounding around 15.94Mb (x-axis). Dark shades indicate high coverage, light shades low coverage. The 10kb intervals used to define source loci are delineated by vertical blue lines. The source locus giving rise to the SV-QTL in (a), (b) is marked with a pink double-arrow. Those founder accessions predicted to carry the reference allele (No-0, Ct-1, Mt-0, Wil-2, Ler-0, Tsu-0, Rsch-4, Kn-0, Zu-0, Hi-0, Ws-0) are in green, those predicted to carry the SV are in grey. Genes are annotated in orange.

We found similar effects on the chromosome 4 QTL for resistance to the fungal pathogen *Albugo laibachii*, isolate Nc14^35^ (Figure 6, **Table S7**). Variation in the number of unpaired reads at 9.50-9.51Mb explains 18.3% of the variance in resistance, and is adjacent to a cluster of Leucine-rich repeat genes, and the genes RPP4^36^, BAL^37^ and RPP5. This locus is rearranged in some Arabidopsis accessions and known to be involved in disease resistance^37^; Figure 6 confirms the founder genomes have complex, polymorphic SVs in this region. Since the resistance QTL is not completely ablated by the SV traits associated with it, additional non-structural variants likely contribute to it.

**Figure 6.**
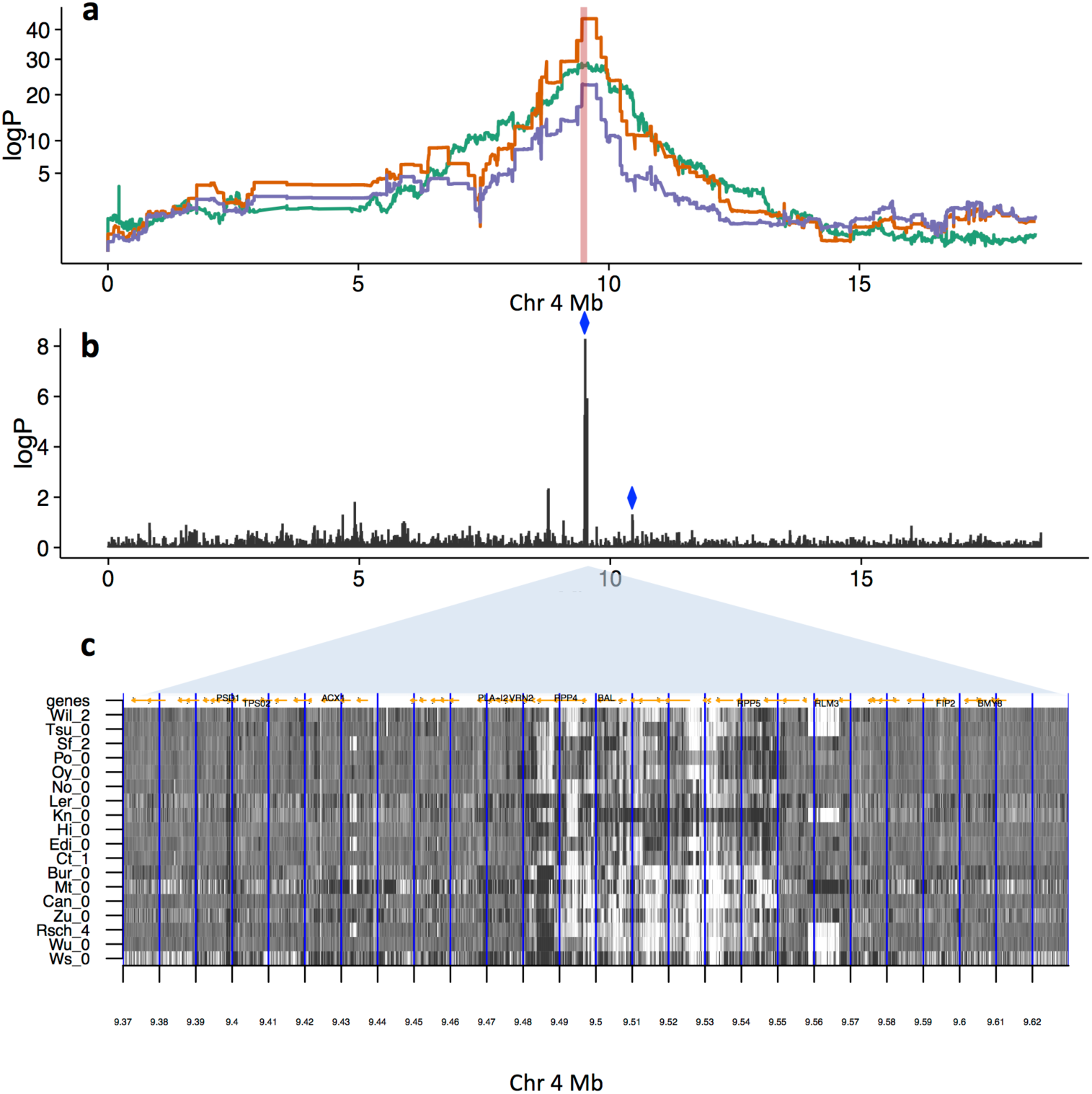
Effects of SVs on resistance to *Albugo laibachii* infection, **(a)** Genome scans on chromosome 4. Orange: Association with resistance. The peak of association for is at 9.50Mb. Green: Association with SV-trait improperly paired reads at source 9.50-9.51. Purple: Resistance after two SV traits have been regressed out measuring improperly paired reads (sources chr4, 9.50-9.51Mb (green line) and chr4, 10.44-10.45Mb (not shown), both marked with blue diamonds in fig (b)) that together explain 24.7% of the phenotypic variance. **(b)** logP of association between SV traits for improperly-paired reads and the resistance trait. There is a cluster of associated traits near 9.50Mb, in addition to the more weakly associated trait at 10.44-10.45Mb. (c) Structural variation in high-coverage sequence in the MAGIC founders around 9.50Mb. Shown is the number of improperly-paired reads (dark: high values, light: low values) in 18 accessions (labeled on y-axis), between 9.37-9.63Mb (x-axis). The 10kb intervals used to define source loci are delineated by vertical blue lines. There is a region of complex structural variation spanning 9.48-9.55Mb approximately, with considerable variation between the founder accessions. Genes are marked by orange arrows, and selected genes, some implicated in disease resistance at this locus, are labelled.

Importantly, Figures 5b,6b show that correlations between SV traits and phenotypes are tightly localized, generally to width of a single SV trait window, in contrast with wider linkage disequilibrium decay seen in QTL genetic mapping (Figure 5a). Consequently correlations between SV traits and physiological traits can sometimes pinpoint causal variants within physiological QTLs which are otherwise too broad to localize (mapping resolution in the MAGIC population is about 200kb^25^).

We also corroborated studies^21,24^ showing SVs associate with gene dysregulation, even when the gene sequence is undisturbed. Within those SVs with mapped breakpoints, 119 genes spanned the breakpoints, 6,909 lay inside the SVs (**Table S8**) and 21,747 outside. Using RNA-seq from 200 MAGIC aerial seedlings, scaled expression variance increased among genes spanning breakpoints (t-test: *P* < 9×10^−3^) and within SVs (*P* < 1×10^−13^) (Figure 7a). Similarly larger fractions of lines had silenced transcripts for genes spanning breakpoints (t-test *P* < 1.2×10^−2^) and within SVs (*P* < 2×10^−52^) (Figure 7b). Expression within SVs was more correlated with local SV traits than outside SVs (F-test *P* < 2.1×10^−6^) (Figure 7c).

**Figure 7.**
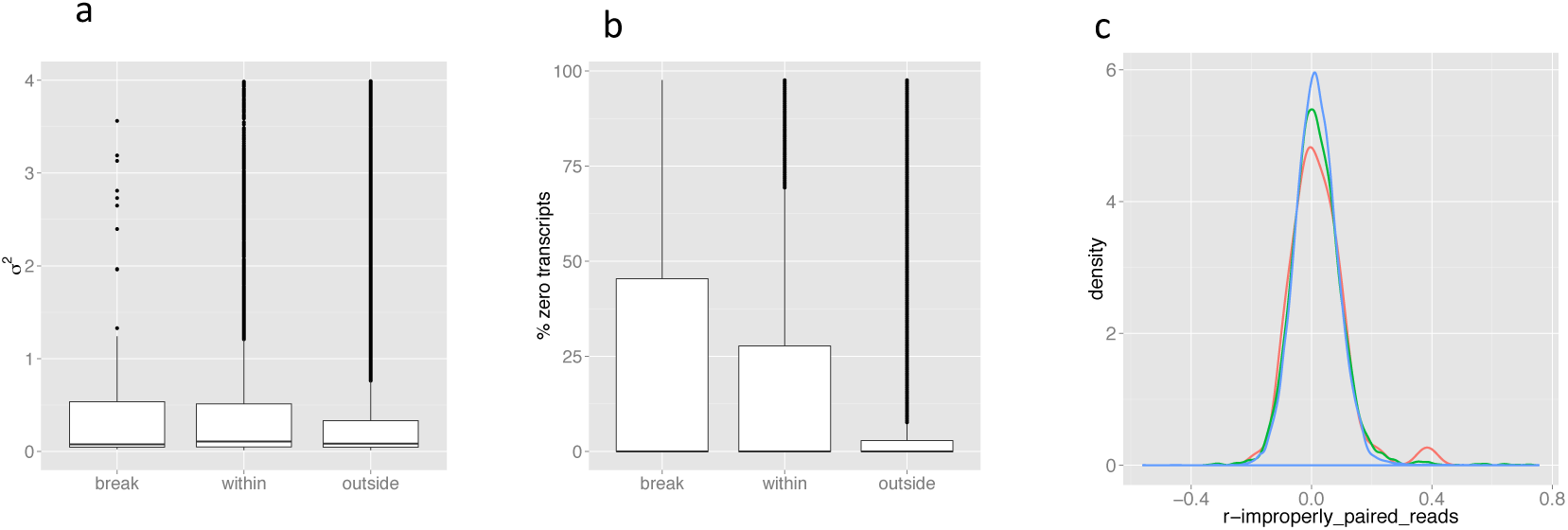
Variation of expression in 200 MAGIC leaf transcriptomes, in genes spanning SV breakpoints, within SVs or outside SVs. **(a)** Boxplots of transcript variance (scaled by the mean). **(b)** Boxplots of the fractions of silenced genes (**c)** Distributions of the Pearson correlations between gene expression and number of abnormal insert size reads in the locus containing the gene (red: spanning breakpoints, green: within SVs, blue: outside SVs).

### Effects of SV-traits on Heritability

Finally, we treated the SV traits as if they were quantitative noisy genotypes to define pair-wise correlations between MAGIC lines, as weighted correlations of their SV traits (**Methods**). We constructed SV genetic relationship matrices (GRMs) *K_SV_*, which we used to compute the SV-heritability 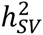 of each of the physiological traits mapped above by analogy with the mixed models used for estimating SNP-based heritability^38^. This idea is similar to the use of gene expression data to model intersample relationships^39^. We also compared these SV-heritabilities with those obtained from “classical” haplotype *K_H_* or SNP-based *K_SNP_* GRMS (Table 2). *K_H_* was computed from the identity between haplotype mosaics (and so measures identity by descent), *K_SNP_* and *K_SV_* were computed from the correlations of 1.2M imputed SNPs or 12k SV-traits respectively (Methods). We also computed SV heritability when only the most variable 50% or 25% of SV-traits were included, to investigate if heritability was concentrated at the most structurally variable loci.

**Table 2.**
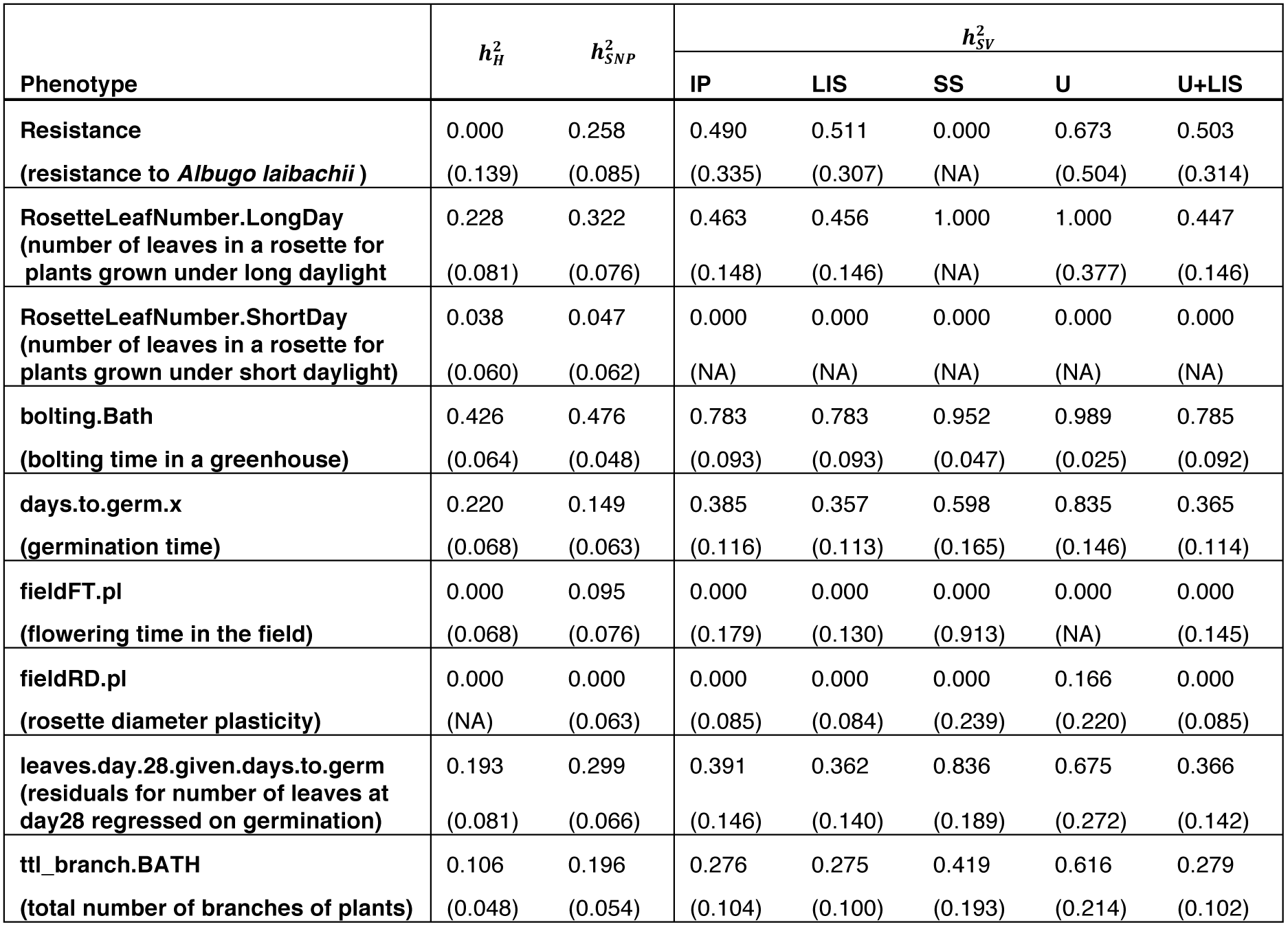
Estimates of heritability. 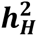 is haplotype-based heritability. 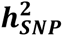 is SNP-based heritability. 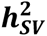 is the heritability estimated from structural variant anomaly traits. Numbers in brackets are the standard errors of the heritability estimates above. ER: Excess Reads, IP: Improperly-paired, LIS: Large Insert Size, SS: Same Strand, U: Unpaired, U+LIS: Unpaired or Large Insert Size. Heritabilities for excess reads are not reported because the fraction of bins in any individual containing non-zero entries was too small.

As expected, SNP-based heritability 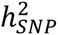 is generally similar to haplotype-based heritability 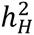 for all phenotypes tested. However, the heritability 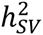 captured by the six measures of SV anomaly is more variable, sometimes being close to zero, but sometimes exceeding classical heritability by a considerable margin (Table 2). The standard error of 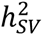 was typically about twice that of 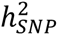 or 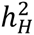, (approximately 0.1 compared to 0.05), presumably reflecting greater uncertainty in SV-traits than in SNPs or haplotypes. Therefore the larger heritability estimates should be treated with caution. Nonetheless, for phenotypes such as times for germination or bolting, the standard errors of all estimates are comparable at ~0.05 and is possible to make meaningful comparisons. Figure 8A,B illustrates likelihood curves the times to germination (A) bolting (B), for SNP, haplotype and large insert-size anomalies. Visualising the entire curves gives a better sense of the uncertainty of the maximum likelihood estimates at the curves’ minima (the standard errors in Table 2 are asymptotic estimates based on the curvature at these minima). The Figure 8B shows that for bolting time the heritability attributable to all largeisize SV-traits, 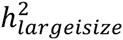, is close to 80%, compared to 40-50% for haplotype or SNP-based estimates. As the fraction of SV traits is reduced by progressively removing those traits with lower variance, 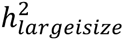 reduces to that of SNPs or haplotypes. This suggests that there is genome-wide structural variation that is not tagged by standard genetic variation, and which has important effects on specific phenotypes. These effects are not universal, as Figure 8A shows for germination time, where heritability is similar for all estimates.

**Figure 8.**
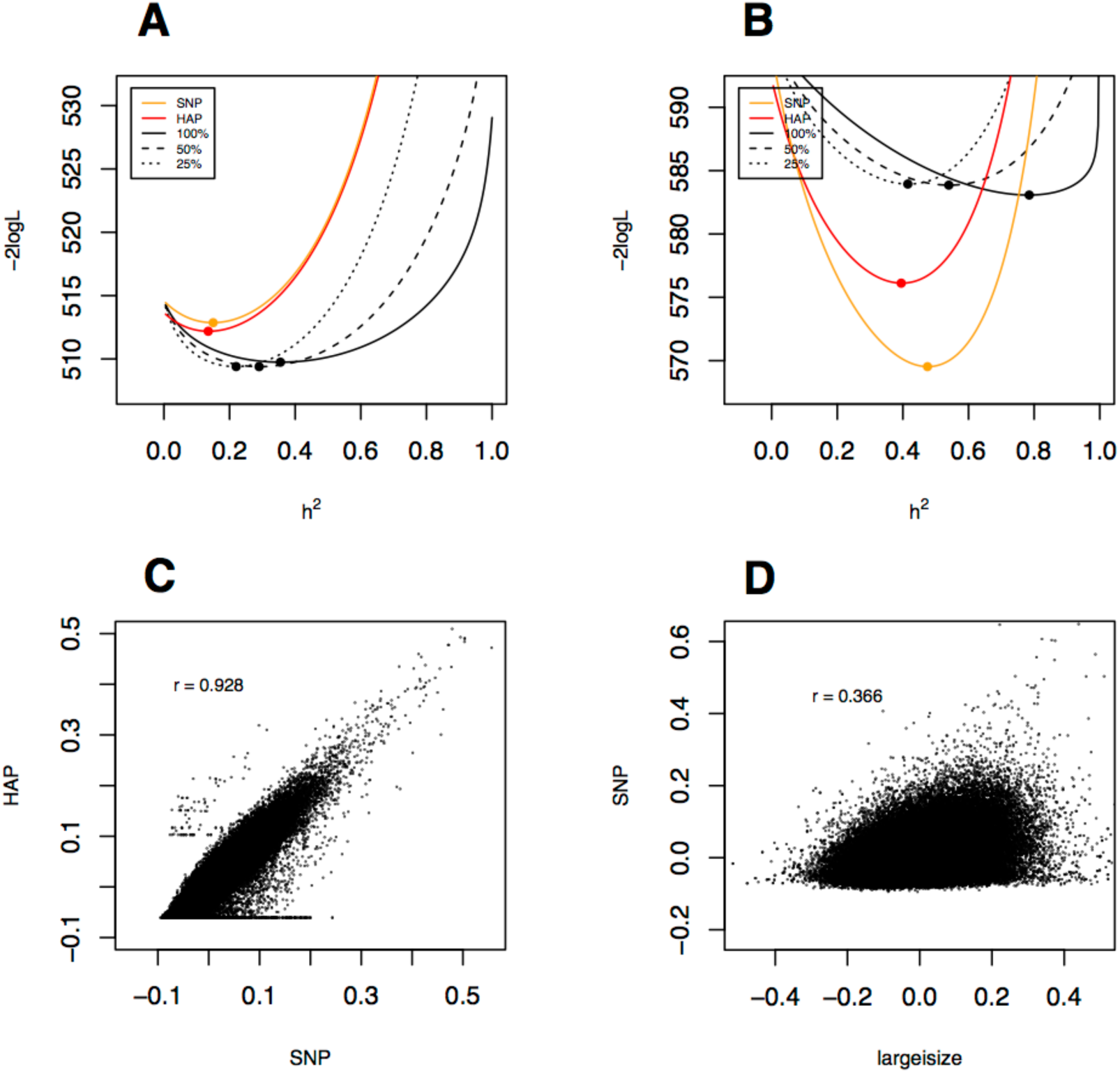
(a,b) log-likelihood curves for two phenotypes bolting.BATH (large insert size read anomalies) and RosetteLeafNumber.ShortDay (unmapped read anomalies), illustrating contrasting behavior of heritability estimates based on structural variants, SNPs and haplotypes. Log-likelihood curves as functions of heritability are plotted for the GRMs estimated from SNPs, haplotypes and various fractions of anomalies. The maximum likelihood estimates of each heritability measure correspond to the minima of the corresponding curves, and are marked with dots. (c,d) Scatter plots comparing the off-diagonal elements of genetic relationship matrices. (c) *K_SNP_* vs *K_H_*; (d) *K_SNP_* VS *K_largeisize_*.

The relative independence of the heritability estimates borne out by low correlations between the corresponding elements of SNP and SV-based GRMs, which range around 0.3 depending on the anomaly type (Figure 8D shows the relationship between GRMs computed from SNPs vs large insertsize anomalies), compared to the correlation of 0.93 between SNP and haplotype based GRMs (Figure 8C).

## Discussion

Our aim has been to understand better the architecture and impact of structural variation in populations sequenced at low coverage. We used a strategy that combines analysis of read-mapping signatures commonly used to detect SVs in individuals sequenced at high coverage, with association mapping in populations^40^. A somewhat related concept was used for mapping un-localised contigs into reference assemblies based on linkage disequilibrium^41^.

In doing so, we have generated a partial catalog of SVs in Arabidopsis, although our purpose is not to call SVs systematically, a task that remains challenging with short reads. Rather, we have shown how SVs’ impact can be assayed without necessarily calling them or mapping their breakpoints. In this way, we can distinguish transpositions from local SVs, and determine the approximate locations of transpositions. The privileged role of the reference genome in the analysis means that some transpositions appear as deletions, so we probably have underestimated their true frequency. Despite this, a quarter of the SVs we detected are transpositions. Given the large numbers of transposable elements in Arabidopsis - over 11,000 from over 300 families are annotated in the reference ^24^ – this is unsurprising. However, many of the SVs we mapped are too large, covering tens of kilobases, to be single transposon-mediated events.

In the minority of cases where we delineated breakpoints exactly, we often found SVs are complex combinations of different SV types. But often breakpoints were not simple cut-and-paste transformations of the reference genome, as illustrated in Figure 6c. Indeed, it is impossible to determine precisely the changes that led to many observed structural variants.

Because we used ultra-low-coverage 0.3x sequence data, we divided the Arabidopsis genome into 10kb bins when counting read-mapping anomalies. With higher coverage and a larger sample size it would be possible to use a larger number of narrower bins, thereby improving resolution. The public release of over 3000 rice genomes sequenced at ~14x ^42^ and over 1000 Arabidopsis accessions sequenced at over ~20x ^43^ means that there are now large collections of inbred plant genomes available for analysis. Both of these sets are worldwide surveys of germplasm, in which we expect SVs to contribute significantly to, and be confounded with, their extensive population structure, in contrast to the MAGIC population used here. Disentangling these effects will be a challenging but important task.

Mapping SVs in a population brings new insights to the problem of QTL analysis. First, an SV trait inside a QTL may entirely explain the genetic effect at the QTL, and hence provide support for being the causal variant (e.g. Figure 6). Second, SV traits are much more tightly localized than are QTLs: there is little or no correlation between neighbouring SV traits so there are no effects of linkage disequilibrium. Our analysis also shows that expression of genes is often dysregulated or even silenced within large SVs, raising the possibility that an SV causes multiple regulatory and phenotypic effects.

Finally we have shown that even in a population like Arabidopsis MAGIC where the local haplotype space is known, structural variation has an impact on heritability that cannot be explained by standard genetic variation. This is unexpected given the breeding history and genetic architecture of the MAGIC lines. For if an SV segregated among the founders of the MAGIC lines, then it should be tagged by the local haplotype context, and therefore contribute to both 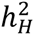 and 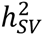.

One possible explanation is that structural variation at loci rich in mobile elements accumulates within each lineage, leading to SVs that are private to each MAGIC line but tend to occur at the same loci, thereby creating similar phenotypic effects. Supporting this, in our analysis the SV-relationship matrix is calculated empirically, without regard to the ancestry of the MAGIC lines, being solely a function of the counts of read-mapping anomalies. Therefore, recalling that the history of each MAGIC line includes a private lineage of at least five generations of selfing, should SVs accumulate recurrently but independently in different lineages, then these could generate phenotypic associations invisible to SNP or haplotype variation. In Arabidopsis, it is known that some mobile elements are methylated, often in response to environmental cues, and that such methylation plays a role in the epigenetic control of certain phenotypes^44^. Testing this hypothesis in Arabidopsis MAGIC lines would require complete and precise reassembly of each genome using long reads, annotation of mobile elements and determination of their methylation status.

The general role that recurrent, but independent, genomic rearrangements might play in Arabidopsis and in other species remains to be seen, but there is no *a priori* reason why it should not be a driver of phenotypic variation. The approach used here may therefore have wider application to other populations, both to characterize the extent of transpositions and the impact of cryptic structural variation on phenotypes.

## Methods

### DNA extraction and sequencing

MAGIC lines were grown at Bath (lab of P. Kover) or Oxford (lab of N.P. Harberd) in greenhouses or growth chambers respectively. Leaves were harvested for DNA extraction. DNA isolation was performed at the John Innes Centre, in 96 well plates using the DNeasy 96 Plant Kit and DNeasy 96 Protocol (www.quiagen.com). Sequencing was performed by the Oxford Genomics Centre.

### Genomic DNA library construction and multiplexing

Samples were quantified using the Quant-iT™ PicoGreen ^®^ dsDNA Kits (Invitrogen) and a Genios plate scanner (Tecan) according to manufacturer specifications. Sample integrity was assessed using 1% agarose gel. Approximately 300ng of DNA were fragmented using a Covaris S2 system with the following settings: Intensity: 5, Duty Cycle: 20, Cycles per Burst: 200, Time: 60 sec. Distribution of fragments after shearing was determined using a Tapestation D1200 system (Agilent/Lab901). DNA Libraries were constructed using the NEBNext DNA Sample Prep Master Mix Set 1 Kit (NEB) with minor modifications and a custom automated protocol on a Biomek FX (Beckman). Ligation of adapters was performed using Illumina Adapters (Multiplexing Sample Preparation Oliogonucleotide Kit). Ligated libraries were size selected using Ampure magnetic beads (Agencourt). Each library was PCR enriched with 25 *µ*M each of the following custom primers:

Multiplex PCR primer 1.0

5’-

AATGATACGGCGACCACCGAGATCTACACTCTTTCCCTACACGACGCTCT

TCCGATCT-3’

Index primer

5’-

CAAGCAGAAGACGGCATACGAGAT[INDEX]CAGTGACTGGAGTTCAGACGTGTGCTCTTCCGATCT-3’

Indexes used were 8bp long (manuscript in preparation). Enrichment and adapter extension of each preparation was obtained using 5*µ*l of size selected library in a 50 *µ*l PCR reaction. After 10 cycles of amplification (cycling conditions as per Illumina recommendations) the reactions were purified with Ampure beads (Agencourt/Beckman). The final size distribution was determined using a Tapestation 1DK system (Agilent/Lab901). The concentrations used to generate the multiplex pool were determined by Picogreen. The library resulting from the pooling was quantified using the Agilent qPCR Library Quantification Kit and a MX3005P instrument (Agilent) before sequencing on an Illumina GAIIx as 50bp or 100bp paired end reads. All steps for library construction, including the setup of the PCR reaction were performed on a Biomek FX (Beckman). Post PCR cleanup was carried out on a Biomek NXp (Beckman) whereas a Biomek 3000 (Beckman) was used to generate the pools of 96 indexed libraries.

### Processing Sequence Reads and SNP Calling

The Illumina reads were mapped to the *A thaliana* reference genome (TAIR10) using Stampy version v1.0.20 ^25^. Alignments were stored in a separate BAM file for each MAGIC line. Previous sequencing for the 18 MAGIC line progenitors had produced a catalogue of 3,316,270 segregating SNPs^17^. We ran GATK v2.6^26^ on the segregating SNPs to call variants for the 19 founders, setting the following read filters: Allele Balance, BaseQualityRankSumTest, Clipping RankSumTest, Coverage, DepthPerAlleleBySample, FisherStrand, GCContent, HaplotypeScore, LowMQ, MappingQualityRankSumTest, MappingQualityZero, MappingQualityZeroBySample, RMSMappingQuality, ReadPosRankSumTest. We filtered out SNPs that were triallelic, within transposons, or heterozygous for any founders.

### Definition of Structural Variant Traits

We divided the TAIR10 (Col-0) reference genome into 11,915 abutting 10 kb segments. Within each segment we computed six measures of anomalously mapped reads that are signatures of SVs. Let *R* be the set of all reads mapped to a genome of length *L*;*ρ* is the number of reads in *R*, and *ρ_l_* the number of reads mapped to a segment *l* of length 10kb. The read anomaly measures computed in each segment are:

1. **High read coverage: *ρ_hc_* = *ρ_l_* – 1.5*E*[*ρ_l_*], where** 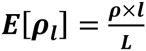 is the expected read coverage of the segment
2. **Unpaired reads: *ρ_u_*** number of reads mapping to the segment whose pair is not mapped
3. **Pairs on the same strand: *ρ_s_*** number of reads with pair on the same strand
4. **Reads with large insert size: *ρ_is_*** number of read pairs with insert size outside the range *m_s_* ± *IQR_s_* or mapped to different chromosomes, *m_s_*, *IQR_s_* being the median and interquartile range of insert sizes of all the reads in the sample.
5. **Unpaired reads or with large insert size: *ρ_ui_* = *ρ_iu_* + *ρ_i_***
6. **Improperly paired reads: *ρ_uis_* = *ρ_u_* + *ρ_i_* + *ρ_s_***

The last two traits are combination of others – certain SV types can cause multiple anomaly signatures, so merging them may increase power. Each type of read pair anomaly was measured in each of the 11,915 10kb segments, determining 71,490 traits in total.

### Genome scan

We treated the SV traits like a gene expression eQTL study, performing a genome scan for each one. Association was tested by fitting trait vectors to the imputed ancestral haplotype at each locus in the 488 genome mosaics. In combination, the mosaics partitioned the genome into 16,700 haplotype blocks, with the ancestral haplotype of all lines unchanged in each one. Let *Y_Ai_* be the number of anomalous reads of a certain type at source segment *A* in line *i*. At every haplotype block *p* we fit the linear model:

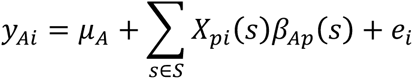

*μ_A_* is the average trait value at *A*, *X_pi_*(*s*) is a binary indicator of whether line *i* carries haplotype *s* at *p*, *β_Ap_*(*S*) is the effect of founder haplotype *s* and *e_i_* the (Normally distributed) error. The founder effects *β_Ap_* (*s*) were estimated by an one-way ANOVA with null hypothesis:

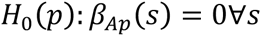

### Genome-wide significance

We denote the location of a sink locus associated with an SV trait an SV-QTL. Each genome scan of a given SV trait returns a p-value *π_Ap_* for each of the 16,700 scanned blocks *p*. We selected as candidate SV-QTLs for each mapped trai the locus with maximum genome-wide negative logarithm of *π_Ap_*, i.e.

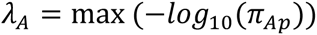

To control for the number of test in each scan (16,700) and correct for associations driven by outliers we performed 100 permutations *T_A_* of the trait vector *y_Ai_* and repeated the mapping for each one. We then fitted a generalized extreme value distribution (GEV), using R evd package on the *λ_A_*(*t*), *t* ∈ *T_A_* values obtained by permutation, from which we obtained a genomewide corrected p-value:

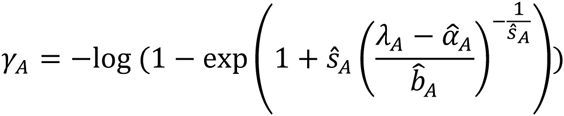

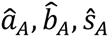 are the MLEs of *a_A_*, *b_A_*, *s_A_*. Study-wide SV-QTLs are selected at *FDR* < 10^−2^ (*γ_A_* < 10^−3^). At this FDR we mapped 10,275 SV-QTLs in total. **Table S3** shows mapped QTLs per read anomaly category. 3,773 SV-QTLs had coincident sources and sinks, probably corresponding to the same SV, and so were counted only once. SVs are tabulated in **Table S4**.

### *Cis* and *trans* SV-QTLs

In total we mapped 6,502 distinct SV-QTLs, each corresponding to a unique SV. SV-QTLs with sources and sinks within 2Mb from each other were classified as *cis*, and the rest as *trans*.

### Prediction of SV allele frequency

We predicted the founder haplotypes carrying SVs at the origination of the population, using the fact that SV haplotypes will have elevated anomalous reads at the source. For each SV-QTL the founders’ contributions were arranged as a 19×19 table *T* whose cells (*i*, *j*) carry the sum of read anomalies (of a certain type) at the source for all lines carrying haplotype *i* at the sink and haplotype *j* at the source. A founder is classified as carrying the SV if its corresponding row has generally higher values than the rest of the table (we note that in *cis* QTLs the matrix is almost diagonal).

For each cell of *T*, let *t_ij_* be the sum of trait values for all genomes carrying haplotypes *i*, *j* at the sink and source, respectively. The contributions of founder *i* are estimated as the ‘’row’’ effect: *r_i_* = Σ*_j_ t_ij_*, which are reorder such that *r*_1_ >…> *r*_19_, with *r*_1_ ≈…≈ *r*_19_ under null. We rejected the null hypothesis if there is a set {*r*_1_,…*r_k_*} such that *r*_1_ >…> *r_k_* > *r*_*k*+1_ >…> *r*_19_. There are 18 such possible sets. The z-score of each set *k* is:

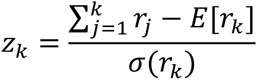

*E* [*r_k_*] and *σ*(*r_k_*) are estimated by 1000 permutations of *T*, denoted as *R_z_k__*. We choose *k* such that *z_k_* is maximised. *k* is significant, hence the corresponding founders carry the SV if the permutation p-value 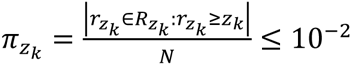. Examples of *cis* and *trans* SV-QTL tables with detectable founders are shown in **Figure S1**. The test predicts founder haplotype groups at 2,391 SV-QTLs.

## Heritability

For a given phenotype (such as germination time) measured in the MAGIC lines, the phenotypic variance matrix is represented by the mixed model 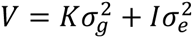 where *k* is the genetic relationship matrix (GRM) and *I* the identity matrix. The phenotypic heritability is

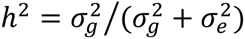

We computed genetic relationship matrices *k* between the inbred MAGIC lines in three ways:

### Identity By Descent (haplotype-based) K_H_

We used the representations of chromosomes as homozygous mosaics of the 19 founders to determine identity by descent (IBD). Across all *N* MAGIC lines, we identified the union of the mosaic breakpoints, and then segmented the genome of each MAGIC line according to these breakpoints. Thus by construction, the founder haplotype for each line is constant within each segment. The founder haplotype in segment *L* in line *i* is represented by an indicator matrix *H_iLf_* which is 1 if the founder is *f* and 0 otherwise. If *W_L_* is the fraction of the genome covered by *L*, and *f_ijL_* = Σ*_f_ H_iLf_ H_iLf_* is the indicator of whether lines *i*, *j* are IBD at *L*, then the fraction of the genome that is IBD for lines *i*, *j* is

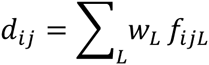

This matrix is then standardised to take the form of a genetic relationship matrix. Let *P_L_* be the probability that, given the observed population-wide founder haplotype fractions at *L*, two randomly-sampled lines are IBD, i.e.

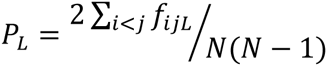

Define

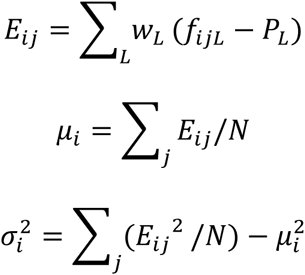

in order to compute the standardised IBD matrix ***K_H_***

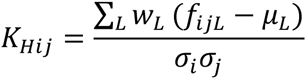

which has main diagonal 1 and off diagonal elements in the range [−1,1].

### Identity by State (SNP-based) K_s_

SNPs were imputed in the MAGIC lines by using the haplotype mosaics and the catalog of variants in the 19 founders. We treated each MAGIC line as being homozygous. We investigated down-sampling the number of SNPs (as the total is over 1 million). Subsamples of between 1% and 10% of the total SNPs were used to define the GRM in the usual way for a homozygous population. Thus if *S_ip_* ∈ {0,1} encodes the SNP genotype in individual *i* and SNP *p*, and if *π_p_* is the allele frequency at *p*, then the normalized genotype is

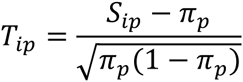

Since the MAGIC lines are almost fully inbred the normalization is different from that in an outbred population under Hardy-Weinberg equilibrium. The SNP-based GRM is the matrix ***K_s_*** with elements

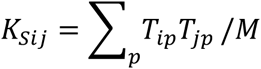

That is,

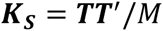

which is always positive semi-definite.

### Read Anomalies K_R_

We constructed read-anomaly GRMs by analogy to SNP-based GRMs. Let *X_iL_* be the read anomaly trait for individual *i* at locus *L*. Let *α_L_* and 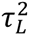 be the sample means and variances:

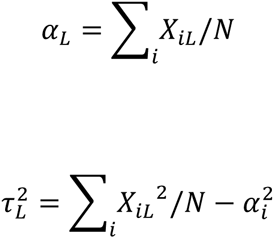

Define the standardized trait matrix *W* with elements

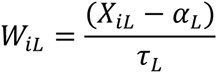

The genetic relationship between individuals *i*, *j* is

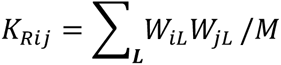

where *M* is the number of loci. Then the read anomaly GRM ***K_R_ = WW'***/*M*.

This formulation guarantees that the GRM is positive. The choice of loci that contribute to the GRM can be varied. Loci at which there is no variation in read anomaly are superfluous and so are ignored. Similarly, loci in which only a small fraction of individuals are anomalous (say <3%) are likely to carry too much weight after normalization and so may optionally be dropped, analogously to the calculation of SNP-based GRMs using only high-frequency SNPs.

In the MAGIC population, each of the 19 founders should be present at a given locus in about 1/19 = 5.5% of lines. Thus an SV that is private to a single founder should give rise to a trait which is null (ie its un-normalised trait value is zero) in 94.5% of lines on average. Loci with a much smaller fraction of non-null trait values might represent private structural mutations.

We computed a separate kinship matrix for each of the six measures of read anomaly, and estimated heritability by maximum likelihood.

### Validation by paired-end data

We used high and low coverage paired-end reads from the 19 founders^8^ and from the MAGIC lines to search for enrichment of read pairs linking the source and sink. For the high-coverage test we restricted attention to the 2,391 SV-QTLs in which founders carrying the SV are predicted, and compared the number of read pairs with one read mapped in the 10kb of the source and within a variably-sized window of *W* kb from the sink (association peak) (*W* ∈ {5,20,30,40,50,100,150,200,400}) in the founders carrying the SV to the remaining founders using two-sided Fisher’s exact tests (FET) at the 5% level of significance. Given the haplotype structure of the MAGIC lines, mapping resolution is variable between QTLs in MAGIC - (200kb being the average in MAGIC^25^) and may depend on the significance of the association. In the low coverage data we performed the same test comparing the 100 lines with the highest read anomaly trait value to the rest of the population.

### Validation by *denovo* contigs

We used BLAT ^45^ to align 5,524,143 short contigs (50-1000bp) from existing *denovo* assembly contigs of the 18 non-reference founder genomes to the reference (Col-0 TAIR10) to identify contigs split across the source and sink locus. After alignment we excluded genomic regions with annotated repeats or transposons and alignments that mapped to over 5 genomic loci. We found 2,619 contigs with alignments split into disjoint pieces over 420 QTLs’ sources and sinks, suggesting a cut-and-paste mechanism. We also found 460,656 (8.3%) shared contigs whose alignments overlapped between source and sink regions (duplications, transposons, Microhomology-Mediated Break-Induced Replication (MMBIR) sites and Non-Allelic Homologous Recombination (NAHR) being possible explanations). We randomized the SV-QTLs by circular genome permutation^46^ to determine whether such split and shared contig alignments are overrepresented near SV-QTLs. In particular, for each SV-QTL *i*. if *a*(*i*), *b*(*i*) are the original position of the source and sink respectively, then a permuted SV-QTL *a_k_*(*i*), *b_k_*(*i*) is defined as:

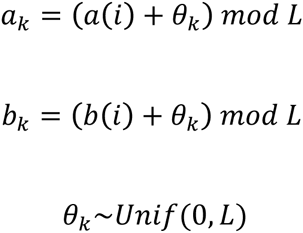

with the requirement that *a_k_*(*i*), *b_k_*(*i*) must be on the same chromosome for *cis* SV-QTLs. We then computed one-sided p-values *π_split_*, *π_shared_*. At the 1% level, *trans* SV-QTLs were enriched for both split and shared alignments and cis only for split.

### Validation by PCR

We designed PCR primers for 77 breakpoints from 44 SV-QTLs predicted from *denovo* contigs. We considered two types of experiments: *type I* experiments had primer oligos corresponding to remote or inverted reference loci so PCR should produce a product in SV genomes and not in the reference; *type II* experiment is a control experiment with the reverse outcome (product in the reference, but not in SV genomes). In total, we designed 96 *type 1* experiments, one for each of the 77 breakpoints, and 19 control (*type 2*) experiments, wherever possible.

We designed 20-30bp primer oligos based on the reference (TAIR10), using Primer3^47^, after masking out repeats, transposons and known polymorphisms. SVs tend to be near such sequence features, so we had to relax the default Primer3 criteria to detect oligos, and in particular we required: (i) Maximum allowed product 1.5kb (ii) Annealing temperature 10-90^o^C (iii) GC-content 10-90% (iv) Self-complementarity 8bp. Primer specificity was tested by BLAT^45^. In 30 (66.6%) SV-QTLs (46 *type 1* experiments) at least one breakpoint was confirmed, i.e. there was at least one *type 1* experiment which amplified in a subset of founder genomes (those carrying the SV) while not producing a product in the reference, as expected. In a further 7 SV-QTLs (15.6%) (15 *type 1* experiments) the founders carrying an SV-QTL amplified successfully, but the reference genome also amplified unexpectedly. This may be due to the presence of highly similar sequence nearby, causing unexpected binding of one of the primers – potentially in the presence of duplications. Indeed in 10 of these experiments evidence of duplications (multiple bands produced by PCR) was detected in more than two founder genomes. However, in all 15 experiments there were at least three founder genomes behaving differently than the reference, indicating that the region is probably structurally variant, although the type of variant may be different to the one predicted by the mapping. We conclude that these results, despite ambiguous, probably indicate SVs, albeit likely polymorphic or of different type than originally predicted. The remaining 16 *type 1* experiments failed amplify in any founder. Of the 19 *type 2* experiments, 16 succeeded (worked as expected), 2 were ambiguous and 1 failed.

In total we confirm at least 30 (66.6%) SV-QTLs with at least one breakpoint, while for a further 7 (15.6%) we have evidence of structural variation – in total up to 82.2% of the tested SV-QTLs are confirmed.

### Association with physiological phenotypes

For each of the six read anomaly categories, we computed Pearson correlations and corresponding p-values between 9 physiological phenotypes and the 11,915 traits measured genome-wide. We selected significant correlations with logP>4. After filtering out correlations driven by outliers (i.e. in which removal of the three most extreme samples reduced the correlation below the significance threshold) we found 549 traits associated with 40 phenotypes. Each physiological phenotype had on average 1.56 associated SV traits of the same anomaly type.

The effect of SVs on each phenotype was measured by a heritability-like measure, 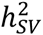, estimated by linear models. Let *y* be the vector of phenotypic values for a physiological phenotype with *k* correlated SV traits (of the same type): *X*_1_,…, *X_k_*, represented by the matrix *X*. The phenotype is modelled as:

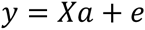

The *k* parameters *a* were estimated using the R function glm() and we computed the residual sum of squares RSS and the total sum of squares (i.e. variance of *y*) TSS. We also computed the individual effect sizes of all traits contributing to the heritability, by fitting simple linear regression models. Based on this analysis SV traits can explain up to 33% of the total phenotypic variance.

We mapped QTLs for the phenotype residuals after regressing all/each associated SV traits of the same type and compared them to the phenotype QTLs.

### Published phenotypes

We used flowering time and rosette diameter data from a field experiment^33^, as well as phenotypes described in previously^25^.

### Phenotyping resistance

Three replicates of each MAGIC line were grown at the University of Bath in 2.5 inch plastic pots. Plants were monitored daily and germination and bolting day recorded. After plants senesced, the inflorescence height and the total number of branches were measured. In a separate experiment, MAGIC lines were grown in growth chambers in P24 plastic trays and sprayed with *A*. *laibachii* race Nc14^35^ when plants were 21 days old. Nc14 zoospores were suspended in water at a concentration of 10^5^ spores per ml and incubated on ice for 30 min. The spore suspension was then sprayed on plants using a spray gun, and plants were incubated in a cold room in the dark overnight. Infected plants were kept in a growth chamber under 10-h light and 14-h dark cycles with a 20°C day and 16°C night temperature^48^. Resistance was defined as absence of pustules on the leaves at 7 days after inoculation. To minimize errors in scoring, resistant plants were monitored up to 14 days after inoculation. The experiment was reproduced twice.

### Collection of RNA

We obtained a subset of MAGIC lines from the Nottingham Arabidopsis Stock Centre (NASC). We grew 209 of the MAGIC lines at 20°C in Percival environmental chambers (Perry, IA, USA) and prepared total RNA as previously described for an earlier study with the MAGIC parental founders^8^; briefly, twenty aerial rosettes from seedlings at the fourth true leaf stage were pooled^8^. RNAseq library construction and sequencing was performed at the Oxford Genomics Centre (Oxford, UK) to produce 2 × 100bp reads using the Illumina non-strand specific method. Per Illumina HiSeq lane, samples were barcoded and run in 13-plexes to give approximately 14 million reads per sample.

### Alignment of RNAseq reads and expression quantification

All libraries were aligned to the TAIR10 reference genome using PALMapper v0.6^49^, following a variation-aware alignment approach. Genome variants collected from the 19 founder strains as well as variants reported for a diverse natural population^28^ were integrated and provided to the aligner as set of known variants. Briefly, the mapper used this set of variants in alignments to prevent reference biases in RNAseq read mapping (see previous work^8^ for a rationale). To facilitate accurate alignments, we further provided splice junction information collected in an earlier study with the founder strains^8^ as well as junction information extracted from the TAIR10 genome annotation. The full alignment parameter set for PALMapper was: -M 3 -G 0 -E 3 -l 12 -L 14 -K 12 -C 14 -I 5000 -NI 1 -SA 5 -UA 50 -CT 50 -JA 15 -JI 1 -z 10 -S-seed-hit-truncate-threshold 100 -report-map-read -report-spliced-read -report-map-region -report-splice-sites 0.9 -filter-max-mismatches 0 -filter-max-gaps 0 -filter-splice-region 5 -min-spliced-segment-len 1 -qpalma-use-map-max-len 10 -f bam -qpalma-prb-offset-fix -junction-remapping <junction_file> -score-annotated-splice-sites <junction_file> -max-dp-deletions 2 -use-variants-editop-filter -use-variants <variant_file> -filter-variants-minuse 1 -merge-variant-source-ids -use-iupac-snp-variants -filter-variants-map-window 20 -iupac-genome -filter-variants-maxlen 100 -index-precache

### Gene expression quantification

We used a custom python script that counted the number of reads overlapping with at least one exonic position of an annotated gene feature. For each read only the best alignment was considered for counting. An alignment was excluded from being counted towards the expression of a gene if (i) at least one position in the alignment overlapped to an annotated intron, (ii) the alignment fell entirely into a region where two or more annotated genes overlap, and (iii) did not start at a position that is part of an exon in all annotated isoforms. For each gene feature the total number of reads passing these filters were used as the expression count.

### Effects of SVs on gene expression

We considered SVs with accurate breakpoints (see **Validation by *denovo* contigs**). 119 TAIR10 genes spanned SV breakpoints (i.e. were disrupted by SVs) and 6,909 were inside them (transposed, inverted or duplicated). Genes were divided into three categories: disrupted by breakpoints, within SV-regions and outside SVs and compared with respect to mean and variance using t-tests. We also computed the correlation of these genes with their local read anomaly values (for the six read anomaly types), i.e. with the 10kb source region that contains the gene and compared the mean and variance (by a t-test and an F-test, respectively) of the Pearson correlation coefficients across categories.

## Acknowledgements

MI, RM were supported by the Wellcome Trust Core Award Grant Number 090532/Z/09/Z. RMC was supported by NSF 0929262. EJO and RG were supported by National Institutes of Health Genetics Training Grant T32 GM07464. We thank Fernando Rabanal for comments on the manuscript.

## Supplementary Figures legends

**Figure S1** Manhattan plots and founder contributions for high read coverage in a *cis* (**a,c**) and a *trans* (**b,d**) SV-QTL. In the manhattan plots the red line shows the source and the association peak the sink of the SV-QTL. In the founder contributions tables rows and columns correspond to founder haplotypes at the sink and source, respectively. The colour hue at each cell is the trait value for each combination of founder haplotypes, darker colour means higher value. In **c** trait values range from 0 to 1000. The figure shows a duplication (confirmed by *denovo c*ontigs) present in 3 founders, namely Bur-0, Col-0 and Edi-0. In **d** trait values (high read coverage) range from 0 to 600 and the figure is showing a *trans* QTL in chromosome 5, present in Bur-0, Oy-0, Po-0, Rsch-4 and Wu-0.

**Figure S2** Distribution of SV allele frequencies, defined as the fraction of founders carrying the SV allele at SV-QTLs.

**Figure S3** Alignment of a 175 kb manually assembled contig from Ler-0, chr5:22.05 – 25.23Mb^31^ to the reference. See legend of Figure 3 for explanation.

**Figure S4** Validation of SV QTL predictions using paired-end data from high coverage sequence in the 19 founder genomes. The figure shows results for 2,391 SV-QTLs for which predictions of founder haplotypes carrying SVs were reliable. Each bar corresponds to a subsample of the SV QTLs with maximum genome-wide logP exceeding a given threshold, with bar height showing the fraction that were supported by reads at P<0.05. The right-most bar shows the results in the entire set of 2,391 QTLs. The test compares the number of reads linking the 10kb source region to a variably sized window around the sink (see **Methods**). The colours are coding the different window lengths used – where different window sizes gave significant results for the same SV QTL we report the smallest.

## Supplementary Table legends

**Table S1** The 19 founder accessions of the MAGIC population of Recombinant Inbred Lines. Shown are the stock centre numbers, the accessions’ names and the place of origin.

**Table S2** The mosaic reconstructions of 488 MAGIC lines. Each row represents one segment of a MAGIC line. **magic**: name of the MAGIC line; **chr**: the chromosome of the segment; **acc**: the founder accession present at this locus; **from.bp, to.bp**: the start and end bp coordinates of the segment (TAIR10); **from.site, to.site**: the start and end position in terms of sites; **len.bp**: the length of the segment in bp; **sites**: the length of the segment in sites; errors: the number of errors (sites whose genotype does not agree with the founder accession genotype); **error.site**: the number of errors divided by the number of sites; **error.bp**: the number of errors divided by the length of the segment

**Table S3:** Catalogue of SV-QTLs detected by genetic mapping of read pair anomalies. Each row represents a distinct SV. SV-QTLs with coincident sources and sinks for different read pair anomalies are merged into a single row. **src.chr, src.pos**: genomic location of the source, defined as the start of the 10kb region in which read pair anomalies were measured; **sink.chr, sink.pos**: genomic location of the sink, defined as the peak of association; **read.anomaly.traits**: read anomaly traits with the sink SV-QTL; **max.gw.logp**: maximum genome-wide logP estimated by the genome scan (*λ_A_*, see **Methods**), fitted.p – extreme value distribution fitted p-value (*γ_A_* - see **Methods**), - **qtl.distance**: *cis* or *trans*; **SV.type**: prediction of the type of structural variant based on the read anomaly traits that had a SV-QTL; **sink.founders** – founder haplotypes predicted to carry the SV, NA means unknown (**Methods**); **known.indel.dist**: distance between the midpoints of the source loci to the nearest large (>100bp) SV from ^17^; **read.support.founder**: P-value of FET comparing read pairs connecting the source and the sink in founders carrying the SV allele from the high-coverage sequencing of the 19 founders (NA if founders are the sink were unknown) ^17^, **read.support.founder.window**: size of window (distance from association peak) containing the significant association in read.support.founder, **read.support.line, read.support.line.window**: same validation test using read pairs from the 488 MAGIC lines; **denovo.contigs**: Boolean variable showing confirmation by at least one *denovo* contig.

**Table S4.** Breakpoints of 420 SVs detected using *denovo* contigs. **founder**: founder genome in which the breakpoint was detected; **source.chr, source.pos**: source position in which the read anomalies were measured; **sink.chr, sink.pos**: SV-QTL position; **source.break.from, source.break.to**: breakpoints detected by *denovo* contigs corresponding to the source area; **sink.break.from, sink.break.to**: breakpoints corresponding to sink; **source.length, sink.length**: length of the structurally variant region in the source and sink regions.

**Table S5.** PCR validation results **(a)** Results per SV-QTL. **QTL_ID**: id of the SV-QTL; **QTL**: coordinates of the tested SV-QTL, position of the source followed by the position of the sink; **confirm**: Y – yes, A-ambiguous, N – no; **dist**: Boolean indicator of whether the SV-QTL is *cis* or *trans*; **read pred**: type of the SV (e.g. transposition, indel etc) predicted by read anomalies, PCR pred: prediction of the type of SV based on the contigs and PCR results. **(b)** Results per experiment. Each row corresponds to a single experiment (unique combination of primers) performed on all 19 founders. **QTL_ID**: ID of the SV-QTL predicted; **type**: type of experiment 1 or 2 (**see Methods); Forward, Reverse**: unique identifier of the primers used, in the form chr_pos_orientation, (e.g. id 4_1853989F means the sequence starting from chr 4, 1853989bp and with forward orientation). The identifier INV means that the sequence has the opposite orientation than expected. The remaining columns correspond to each of the founder genomes, 1 meaning that the experiment amplified, 0 that it did not, and >1 that it produced multiple bands. (**c)** Primer sequences.

**Table S6** Nine physiological traits measured in MAGIC lines that were used in this study.

**Table S7. Effects of SVs on** physiological phenotypes. **Phenotype**: physiological phenotype; **SV trait type**: type of read pair anomaly, **SV-QTL**: number of SV traits that have SV-QTLs, **trans**: number of trans SV-QTLs; **max.source**: position of the source of the maximum-contributing SV trait; **max.sink**: position of the sink QTL of the maximum-contributing SV, or NA if it is not mapped, **physio.QTL**: position of the QTL of the physiological phenotype; **overlap.source, overlap.sink**: Boolean indicator of whether the source or sink of the SV trait overlaps (within 200kb) with the QTL of the physiological phenotype, NA if there is no QTL.

**Table S8**. Association of SVs with gene expression. **gene** – gene id, **chrom, start, end** – gene coordinates, Same QTL: position of the gene relative to SVs (break: the gene is spanning an SV breakpoint, within: the gene is between SV breakpoints so it may be transposed, inverted or duplicated, outside: the gene lies outside SVs), **mean**: mean gene expression, **var**: variance of gene expression, **zeroes**: proportion of silenced (zero expression) transcripts, **r_unmapped_largeisize, r_impop_paired_LT, r_same_strand, r_largeisize, r_excess_reads, r_unmapped**: Pearson correlation coefficients between expression levels and values of the local read anomaly trait, for each of the six anomaly types.

